# Hypothalamic transcriptome analysis reveals the neuroendocrine mechanisms in controlling broodiness of Muscovy duck (*Cairina moschata*)

**DOI:** 10.1101/453241

**Authors:** Pengfei Ye, Min Li, Wang Liao, Kai Ge, Sihua Jin, Cheng Zhang, Xingyong Chen, Zhaoyu Geng

## Abstract

Broodiness, one of the maternal behaviors and instincts for natural breeding in birds, is an interesting topic in reproductive biology. Broodiness in poultry is characterized by persistent nesting, usually associated with cessation of egg laying. The study of avian broodiness is essential for bird conservation breeding and commercial poultry industry. In this study, we examined the hypothalamus transcriptome of Muscovy duck in three reproductive stages, including egg-laying anaphase (LA), brooding prophase (BP) and brooding metaphase (BM). Differences in gene expression during the transition from egg-laying to broodiness were examined, and 155, 379, 292 differently expressed genes (DEGs) were obtained by pairwise comparisons of LA-vs-BP, LA-vs-BM and BP-vs-BM, respectively (fold change≥ 1.5, P < 0.05). Gene Ontology Term (GO) enrichment analysis suggested a possible role of oxidative stress in the hypothalamus might invoke reproductive costs that potentially change genes expression. KEGG analysis revealed glutamatergic synapse, dopaminergic synapse, serotonergic synapse and GABAergic synapse pathway were significantly enriched, and regulator genes were identified. Eight gene expression patterns were illustrated by trend analysis and further clustered into three clusters. Additional six hub genes were identified through combining trend analysis and protein-protein interaction (PPI) analysis. Our results suggested that the cyclical mechanisms of reproductive function conversion include effects of oxidative stress, biosynthesis of neurotransmitters or their receptors, and interactions between glucocorticoids and thyroid hormones and regulatory genes. These candidate genes and biological pathways may be used as targets for artificial manipulation and marker-assisted breeding in the reproductive behavior.

## Introduction

The domestic Muscovy duck (*Cairina moschata*) is an economically important poultry around the world for its unique meat taste and low-caloric content. Almost all varieties of domesticated ducks are descended from the mallard (*Anas platyrhynchos*), apart from the Muscovy duck[1]. However, Muscovy duck has a long incubation period and strong broodiness, which has led to a decrease in egg production and limited the development of the Muscovy duck industry. Thus, broodiness traits in this species should be the primary concern for Muscovy duck industry. Broodiness of the Muscovy duck is characterized by persistent nesting, neck curled backward, clucking and nest defense[2]. The stable laying-broodiness cycles and easily identifiable characteristics of broodiness make this duck species an ideal model to study broodiness.

Reproductive process is strictly controlled by the hypothalamic-pituitary-gonadal axis to coordinate the behavior across reproductive stages[3]. The transition from the egg-laying to brooding is a complex process result of ovarian and oviduct regression, hyperprolactinemia while the termination of ovulation reduces the egg production [4]. The neuroendocrine basis of incubation behavior has been studied in a few domestic species, including the domestic fowl [5], turkeys [6], doves [7] and geese [8]. It is believed that the altered levels of reproductive endocrine hormones, including growth hormone (GH), prolactin (PRL), luteinizing hormone (LH), progesterone (P4), and estradiol (E2), are the major factors inducing the occurrence of broodiness [3-9]. A wealth of information has confirmed the onset of broodiness in poultry is marked by increased hypothalamic serotonin (5-HT), dopamine (DA) and vasoactive intestinal polypeptide (VIP) [9,10]. The phenotypic and physiological factors of broodiness have been extensively studied, but the molecular regulation mechanism of this behavior remains unclear.

In recent years, the development of next-generation sequencing (NGS), has provided a powerful, highly reproducible and cost-efficient tool for transcriptomic research[11,12]. Previous transcriptome analysis of follicles and pituitary revealed the important mechanisms of hormones, autophagy and oxidation in regulating broodiness and egg-laying[13–16]. Few studies on hypothalamic transcriptome analysis of broodiness behavior in Muscovy duck have been reported until now. The activity of egg-laying and broodiness in poultry is usually coordinated with periodically recurring events in the environment and body, such as photoperiod and development of ovary[17,18]. The hypothalamus integrates signals from the environment and body to synchronize the expression of the reproductive behavior through appropriate mediate endocrine, circadian rhythms, ingestion, body temperature and development of gonads[19,20]. Further efforts, including the study of the temporal gene expression profile of the hypothalamus, are essential to elucidate the molecular mechanisms of reproductive behavior in the poultry.

In this study, we systematically examined the dynamics of the Muscovy duck hypothalamic transcriptome at egg-laying and brooding stages and focused on the gene expression differences between the three reproductive stages. Real-time quantitative PCR analysis was carried out to verify the transcriptome results. The purpose of this study was to understand the genetic basis of the transition between the laying stage and the brooding stage at the genomic level, and to lay the foundation for molecular-assisted selection breeding of Muscovy duck.

## Materials and Methods

### Ethics statement

All experimental procedures and sample collection were performed according to the Regulations for the Administration of Affairs Concerning Experimental Animals (Ministry of Science and Technology, China, revised in June 2004) and approved by the Institutional Animal Care and Use Committee of the College of Animal Science and Technology, Anhui Agricultural University, Hefei, China (IACUC No. AHAU20101025).

### Animals and Sample Preparation

The female Muscovy ducks used in this study were obtained from Anqing Yongqiang Agricultural Science and Technology Stock Co., Ltd. China, and were raised according to the standard procedure. There is great variance of reproductive performance and broodiness between Muscovy duck individuals. At 42 weeks of age, individuals with high fertility are in the laying period, and those with low fertility are in the brooding stage. Ten ducks from egg-laying anaphase (LA), brooding prophase (BP) and brooding metaphase (BM) stage were randomly chosen at 42 weeks of age according to their behavior, respectively[3,9]. After euthanized by exsanguination under anesthesia, hypothalamus and ovary samples were collected. The hypothalamus were wrapped in freezing tube, frozen in liquid nitrogen, and then stored at −80°C until further study. The morphologic characteristics of ovary was used to further evaluate individual physiological stage of Muscovy duck. Four hypothalamic samples in each group were randomly chosen for transcriptome study.

### Library construction and sequencing

After total RNA was extracted, eukaryotic mRNA was enriched by Oligo (dT) beads, while prokaryotic mRNA was enriched by removing rRNA by Ribo-ZeroTM Magnetic Kit (Epicentre). Then the enriched mRNA was fragmented into short fragments using fragmentation buffer and reverse transcripted into cDNA with random primers. Second-strand cDNA were synthesized by DNA polymerase I, RNase H, dNTP and buffer. Then the cDNA fragments were purified with QiaQuick PCR extraction kit, end repaired, poly(A) added, and ligated to Illumina sequencing adapters. The ligation products were size selected by agarose gel electrophoresis, PCR amplified, and sequenced on the Illumina Hi Seq™ 4000 sequencing platform generating paired end reads of 150bp each.

### Alignment with reference transcriptome and transcriptome analysis

All the sequences filtered adaptor sequences and low-quality sequences were mapped to the reference transcriptome using short reads alignment tool Bowtie2[21] by default parameters, and mapping ratio was calculated. The Transcriptome Shotgun Assembly project has been deposited at DDBJ/EMBL/GenBank under the accession number GGZN00000000, which used for reference transcriptome in this study. Gene expression was calculated and normalized to RPKM (Reads Per kb per Million reads)[22]. The formula is as follows: RPKM = (1000000*C)/(N*L/1000). To reveal the relationship of the samples, principal component analysis (PCA) was performed with R package models (http://www.r-project.org/) in this experience. To identify differentially expressed genes across groups, the edgeR package (http://www.r-project.org/) was used. We identified genes with a fold change ≥ 1.5,P <0.05 and RPKM ≥ 1in a comparison as significant DEGs[23]. DEGs were then subjected to enrichment analysis of GO functions and KEGG pathways. Trend analysis was performed by Short Time-series Expression Miner software (STEM)[24]. The proteinprotein interaction (PPI) analysis was performed using Cytoscape_v3.2.1. The PPI information was gained with default confidence cutoff of 400 from STRING (https://string-db.org/).

### qRT-PCR analysis of key genes

Gene expression profiles were determined by qRT-PCR on a CFX96 qRT-PCR detection system (Bio-Rad, Hercules, CA, USA) with associated software. cDNA reverse-transcribed from total RNA was used as the template along with SuperReal PreMix Plus with SYBR Green (Tiangen Biotech, Beijing, China) and gene-specific primers designed with Primer Premier 6 software (Premier Biosoft, Palo Alto, CA, USA). Relative abundance of the transcripts was calculated by the comparative cycle threshold method with Beta-actin used as an endogenous control. Reactions were prepared in triplicate for each sample. Relative expression levels were calculated by the 2-ΔΔct method. Variance was performed using t-test in SPSS 22.0 for Windows (IBM, Chicago, IL, USA) and the significance was determined when P < 0.05.

## Result

### RNA sequencing of Muscovy duck hypothalamus

A total of 57,454,328 to 75,309,536 raw reads were generated in each library. After removing reads containing adapters, reads containing poly-N and low-quality reads from raw data, 56,417,422 to 73,275,898 valid reads were filtered out, with the valid ratios being more than 97% in 12 libraries (Table 1). The data quality was good, with Q20 (base sequencing error probability < 1%) > 97% and Q30 (base sequencing error probability < 0.1%) > 94% in each library (Table 1). All read data are available at the NCBI SRA database using the project ID: PRJNA454042. To identify the genes corresponding to clean reads in each library, the clean reads were mapped to the assembled reference transcriptome and mapped reads were counted RPKM (Reads per Kb per Million mapped reads). The mapping results showed that 82.24% to 83.98% of the reads and 71.73% to 83.15% of the genes from each library were perfectly matched to the reference genome. In comparison, 45099297(80.25%) to 55208603(81.94%) in unique mapped reads, and 1024299(1.91%) to 1095515(2.13%) in multiple mapped reads were matched (S1 Table). Principal component analysis (PCA) was performed using gene expression profiles of 12 samples, sample Correlations mapped onto first two principal components. The two principle components explained 99.4% of the cumulative variance, the PC1 explained 88.9% of the variance and the PC2 explained 9.5% of the variance (Fig 1). The results showed that the sequencing quality complied with the requirements of further analyses and gene expression of four biological replicates had the consistent trend.

**Table 1.** **Characteristics of the sequence reads from 12 hypothalamic libraries in Muscovy duck**.

| Sample | Before Filter |  | After Filter |  |  |  | GC<sup>d</sup> |
| --- | --- | --- | --- | --- | --- | --- | --- |
|  | Reads Num | Before Filter Data(G) | Reads Num (%) | After Filter Data(G) | Q20(%)<sup>b</sup> | Q30(%)<sup>c</sup> |  |
| LA1<sup>a</sup> | 62284400 | 8.70 | 60697720 (97.45%) | 8.44 | 97.87 | 94.44 | 0.4981 |
| LA2 | 68574488 | 9.58 | 66688082 (97.25%) | 9.28 | 97.74 | 94.16 | 0.5018 |
| LA3 | 65867842 | 9.20 | 64170830 (97.42%) | 8.92 | 97.8 | 94.23 | 0.5099 |
| LA4 | 62586652 | 8.74 | 60823958 (97.18%) | 8.47 | 97.76 | 94.19 | 0.5014 |
| BP1<sup>a</sup> | 64059674 | 8.95 | 62579376 (97.69%) | 8.70 | 97.91 | 94.5 | 0.504 |
| BP2 | 60205434 | 8.41 | 58590972 (97.32%) | 8.15 | 97.78 | 94.17 | 0.513 |
| BP3 | 58848768 | 8.22 | 57331926 (97.42%) | 7.98 | 97.91 | 94.5 | 0.4995 |
| BP4 | 61416872 | 8.58 | 59805620 (97.38%) | 8.32 | 97.86 | 94.38 | 0.4992 |
| BM1<sup>a</sup> | 57454328 | 8.03 | 56417422 (98.2%) | 7.82 | 98.02 | 94.69 | 0.5177 |
| BM2 | 75309536 | 10.52 | 73275898 (97.3%) | 10.19 | 97.72 | 94.01 | 0.5099 |
| BM3 | 58441114 | 8.16 | 56848862 (97.28%) | 7.91 | 97.84 | 94.34 | 0.5019 |
| BM4 | 67517684 | 9.43 | 65957334 (97.69%) | 9.17 | 98 | 94.71 | 0.5085 |

<sup>a</sup>LA, BP and BM represent egg-laying anaphase, brooding prophase and brooding metaphase, respectively;
<sup>b</sup>Q20 represents the proportion of read bases whose error rate is less than 1%;
<sup>c</sup>Q30 represents the proportion of read bases whose error rate is less than 0.1%;
<sup>d</sup>GC represents guanine-cytosine content.

**Fig 1.**
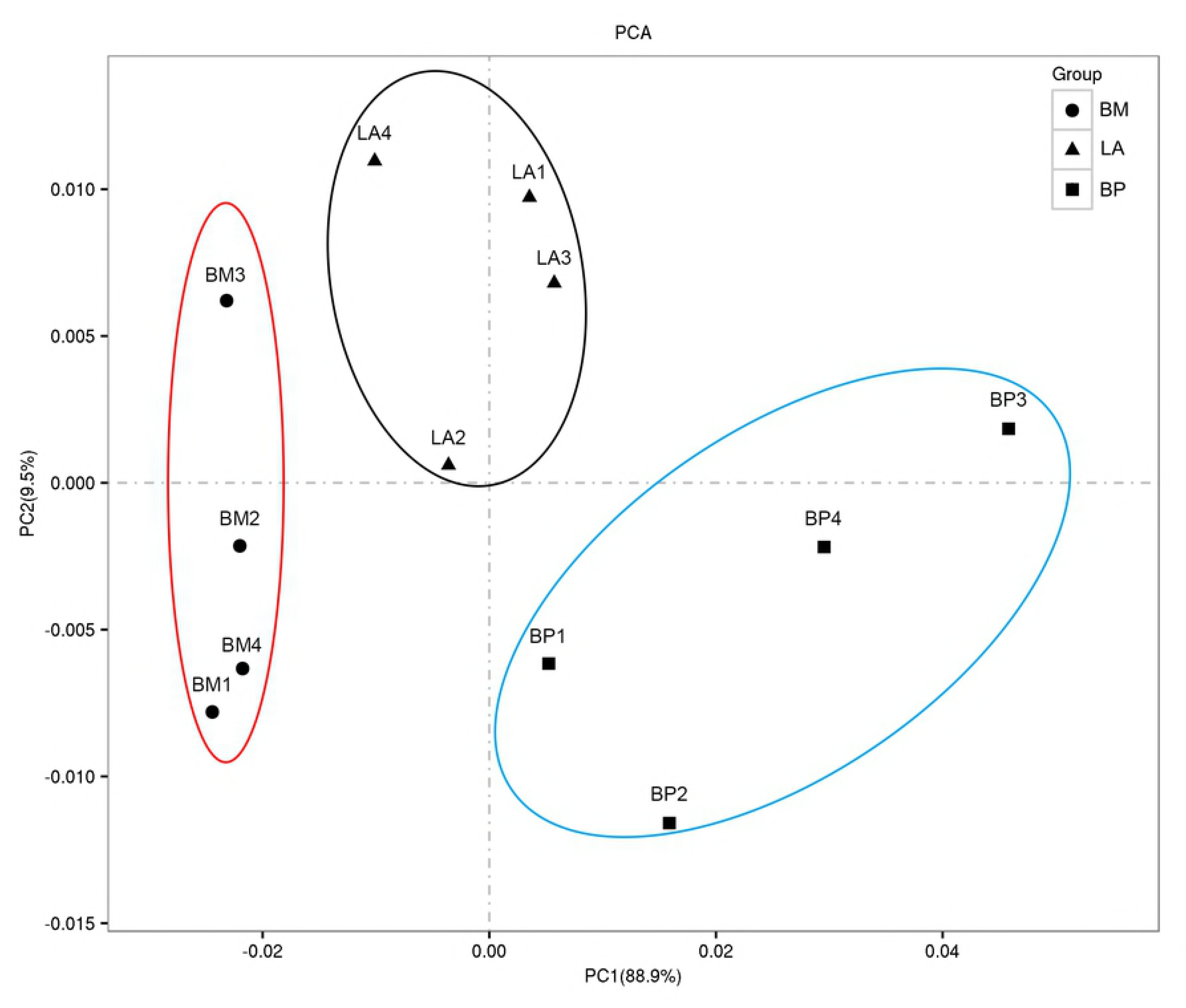
**Principal component analysis (PCA) of sample relationship**. Correlations between 12 hypothalamic sample mapped onto first two principal components of PCA analysis based on genome-wide expression profiles. Triangle nodes standard for sample of egg-laying anaphase (LA), square nodes for sample of brooding prophase (BP), circle nodes for sample of brooding metaphase (BM).

### Analysis of differential expression genes (DEGs)

A cutoff value of 1.0 RPKM was established to evaluate the relative abundance levels of transcripts of the hypothalamus samples that were considered for further analysis. Differences in gene expression during transition from egg-laying to broodiness were examined, and 155, 379, 292 DEGs were obtained by pairwise comparisons of LA-vs-BP, LA-vs-BM and BP-vs-BM, respectively (fold change≥1.5, P < 0.05). Among of them, there were 65/90 (LA-BP), 321/58 (LA-BM), and 243/49 (BP-BM) DEGs with decreased/increased expression level, respectively (Fig 2I, S2, S3 and S4 Tables). Thirty-four DEGs were shared by all 3 reproductive phases (Fig 2II). Differently expressed transcription factors were examined, and 5, 18, 17 TFs were obtained from comparisons of LA-vs-BP, LA-vs-BM and BP-vs-BM with 9 TFs were shared by more than one comparison (S5 Table). The expression levels of 24 randomly selected genes were confirmed by RT-qPCR which consistent with the sequencing results (Fig 3). The log2 ratios of gene expression calculated from RNA-Seq and RT-qPCR were significantly related in LA vs BP, LA vs BM and BP vs BM. Correlation coefficients are respectively equal to 0.7739, 0.9253, 0.9649, with p < 0.0001. Primer sequences for qPCR were shown in S6 Table.

**Fig 2.**
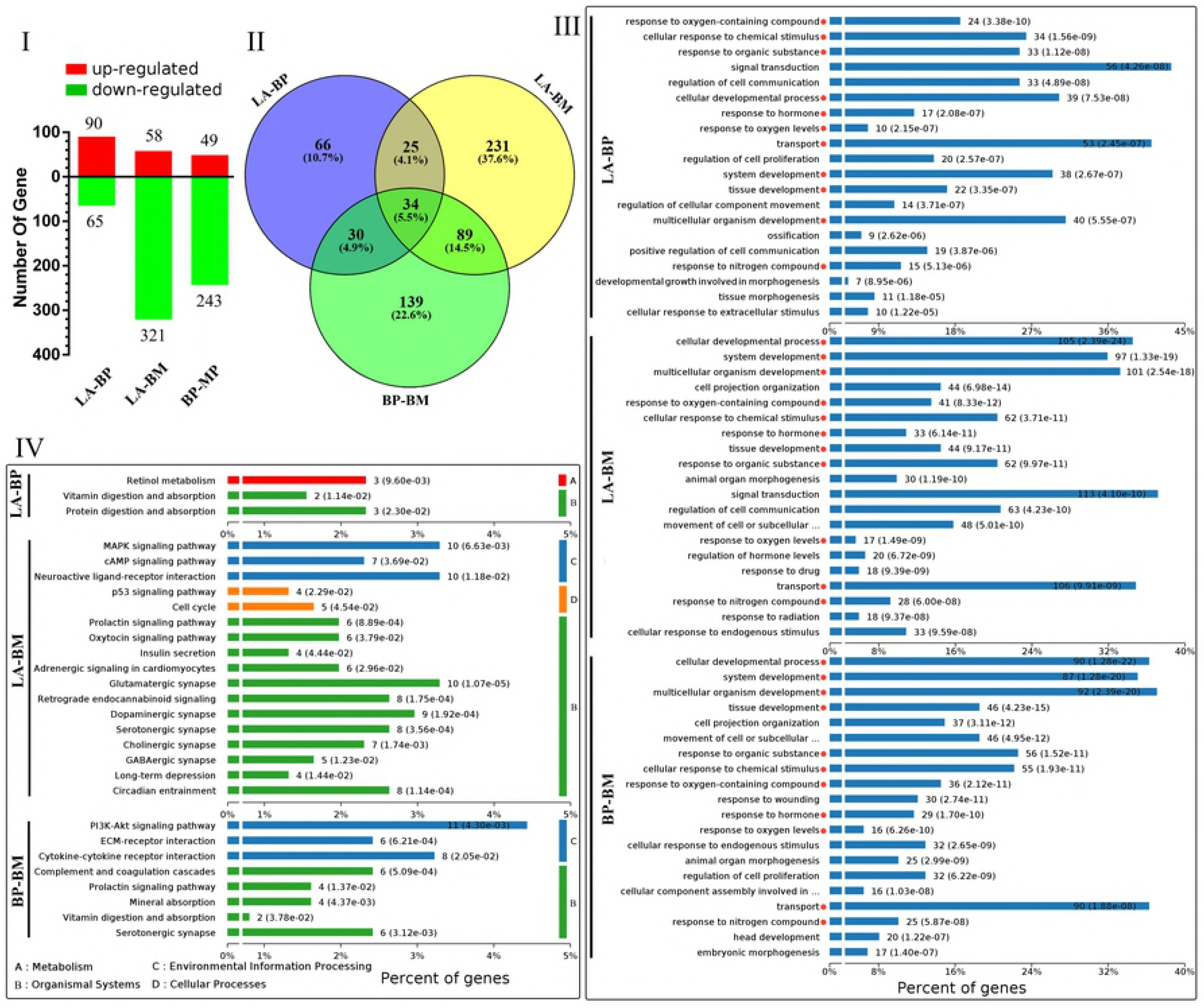
**Statistical results, GO enrichment and KEGG enrichment of different expressed genes (DEGs)**. (I) Histogram of DEGs in pairwise comparisons of three group. Red bar for up-regulated genes, green bar for down-regulated genes. (II) Venn diagrams of DEGs in pairwise comparisons of three group. (III) Top 20 significantly enriched Biological processes of GO enrichment of DEGs in three comparisons. (IV) The significantly enriched KEGG pathways of DEGs in three comparisons.

**Fig 3.**
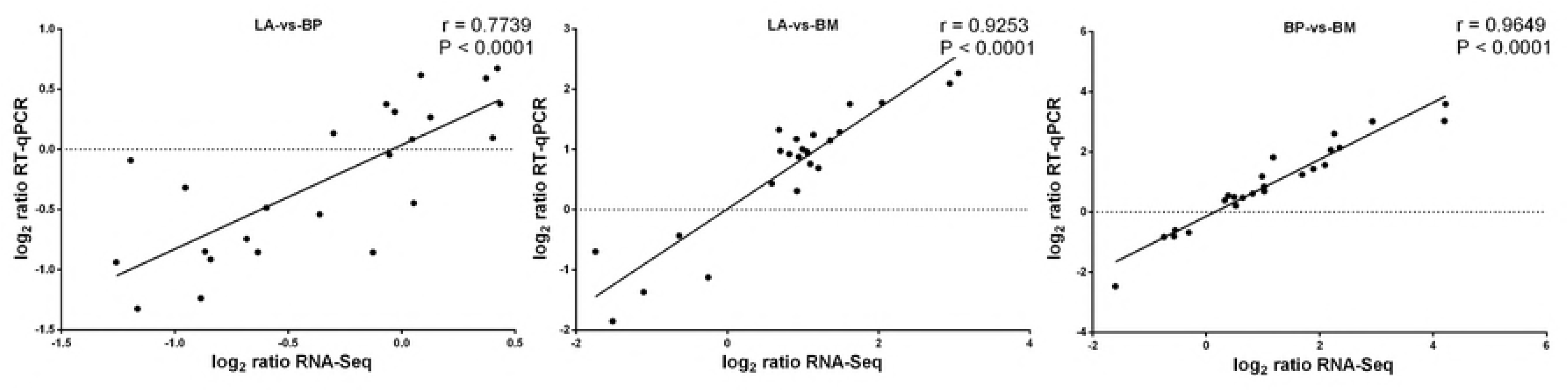
**Validation of the gene expression profile by qPCR**. Plot of gene expression log2 ratios (LA vs BP, LA vs BM and BP vs BM) determined by the RNA-Seq (X-axis) and RT-qPCR (Y-axis) for 24 selected genes (S6 Table). Correlation between RNA-Seq and RT-qPCR was calculated by Pearson product moment correlation in pairwise comparisons of three stages. Each dot represents one tested gene and the plots present linear regression lines, P values and correlation coefficients (r).

### GO and pathway analysis of the DEGs

To determine the function of DEGs during the reproduction process, we performed studies of enrichment of DEGs in GO and KEGG functional categories in pairwise comparisons of LA-vs-BP, LA-vs-BM and BP-vs-BM. Three group DEGs were respective categorized into the 3 main categories of GO classification, including biological processes, cellular components, and molecular functions (S7 Table), and the top 20 significantly enriched biological processes were shown in Fig 2III. In the top 20 biological process, 11 GO terms with red dot behind were both enriched in three libraries, 4 GO terms ‘animal organ morphogenesis’, ‘cell projection organization’, ‘cellular response to endogenous stimulus’, ‘movement of cell or subcellular component’ were significantly enriched in LA-vs-BM and BP-vs-BM. 2 GO terms ‘regulation of cell communication’, ‘signal transduction’ were enriched in LA-vs-BM and LA-vs-BP, of which ‘response to stimulus’ accounted for the highest proportion. Only one term ‘Regulation of cell proliferation’ was enriched both in LA-vs-BP and BP-vs-BM (Fig 2III).

In three pairwise comparisons categories,155, 379 and 292 DEGs were significantly mapped to 3, 17 and 8 KEGG pathways, respectively. The identified pathways were involved in 4 major functional categories (Fig 2IV), showing: (A) Metabolism; (B) Organismal systems; (C) Environmental information processing; and (D) Cellular process. Three KEGG pathways ‘Serotonergic synapse‘, ‘Prolactin signaling pathway’ and ‘Vitamin digestion and absorption’ were enriched in more than one outcome of pairwise comparison. The highest proportion of DEGs were accounted for with nine KEGG terms: ‘Glutamatergic synapse’, ‘Circadian entrainment’, ‘Retrograde endocannabinoid signaling’, ‘Dopaminergic synapse’, ‘Serotonergic synapse’, ‘MAPK signaling pathway’, ‘Neuroactive ligand-receptor interaction’, ‘PI3K-Akt signaling pathway’ and ‘Cytokine-cytokine receptor interaction’ (Fig 2IV, S8 Table).

### Trend analysis

To understand the expression patterns of DEGs, trend analyses of genes showing stage-specific expression were performed and the results displayed in a heat map (Fig 4A). Subsequently, the expression data from LA, BP and BM was clustered into eight profiles by Short Time-series Expression Miner software (STEM), in which 614 DEGs were clustered. STEM analysis revealed that profile 4, 6, 7 clustered into Cluster 1 representing a general upregulation over time. The profile 0, 1, 3 clustered into Cluster that were largely downregulated over time, while profile 2 and 5 contained in Cluster with expression pattern that only changed in BP stage (Fig 4B).

**Fig 4.**
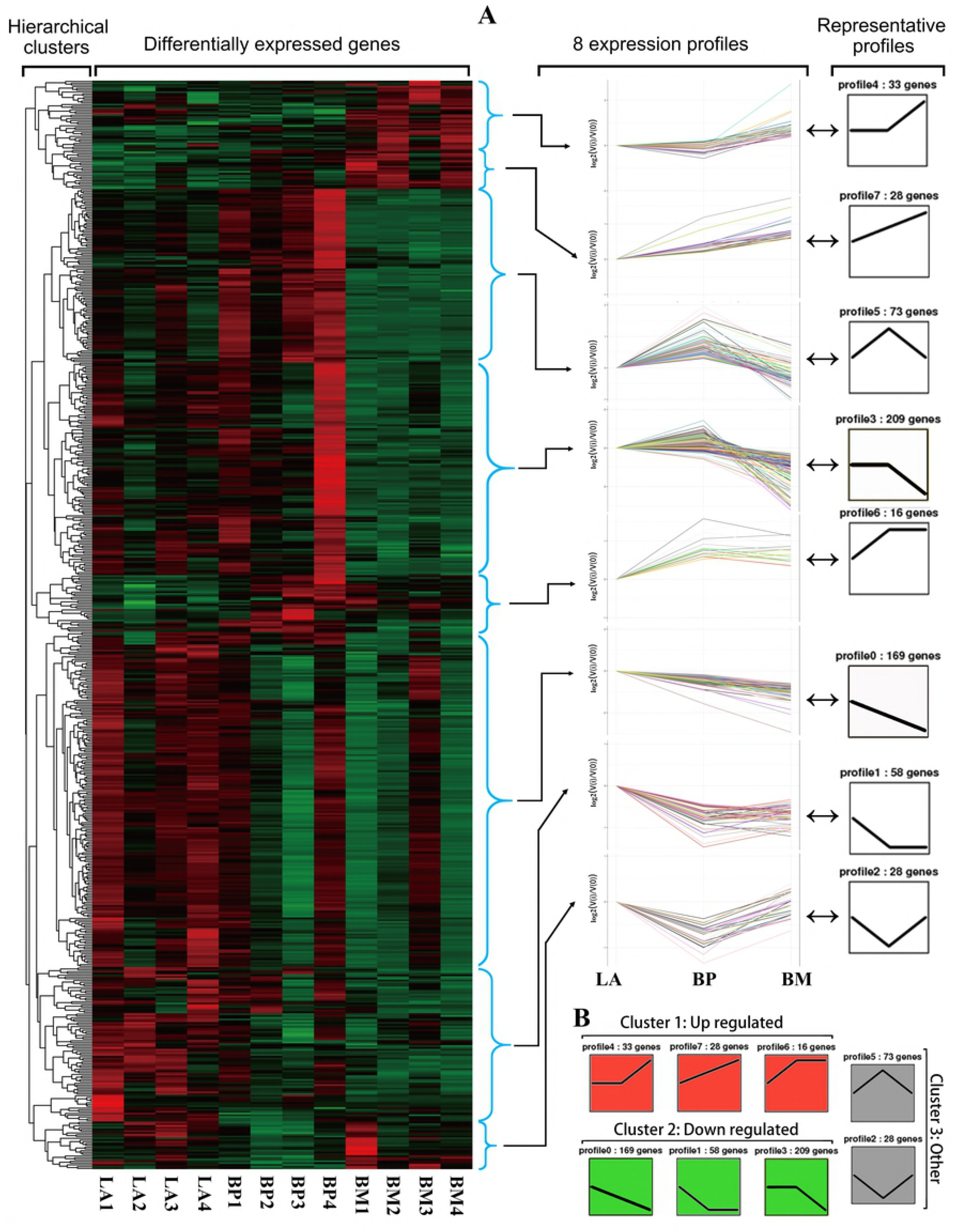
**Overall patterns of differentially expressed genes and the representative profiles in reproductive stages of LA, BP and BM**. (A) Patters were plotted on the heatmap using Treeview. Red represents up-regulated genes while green represents down-regulated genes. Hierarchical clustering is shown on the left. All 8 expression profiles identified by Short Time-series Expression analysis are shown in the middle and the summarized representative profiles are shown on the right. (B) STEM analysis grouping of significant differentially-expressed genes into three temporal profile clusters.

### Protein-Protein interaction analysis

To further extract relevant information from the identified transcriptome data, a more comprehensive bioinformatics analysis of protein-protein interaction (PPI) networks of DEGs was performed (Fig 5). The following network model is generated with cytoscape based on information gained up to 4 level of functional analysis: fold change of genes, protein-protein interaction, KEGG pathway and biological process enrichment. The PPI network of DEGs consisted of 303 nodes and 516 edges and 11 significantly enriched pathways/biological process. The top highest degree genes included *GRP* (Gastrin-releasing peptide)*, FKBP5* (immunophilin)*, TTR* (transthyretin), DIO2 (type 2 deiodinase) and transcription factors of *c-Jun and c-FOS*.

**Fig 5.**
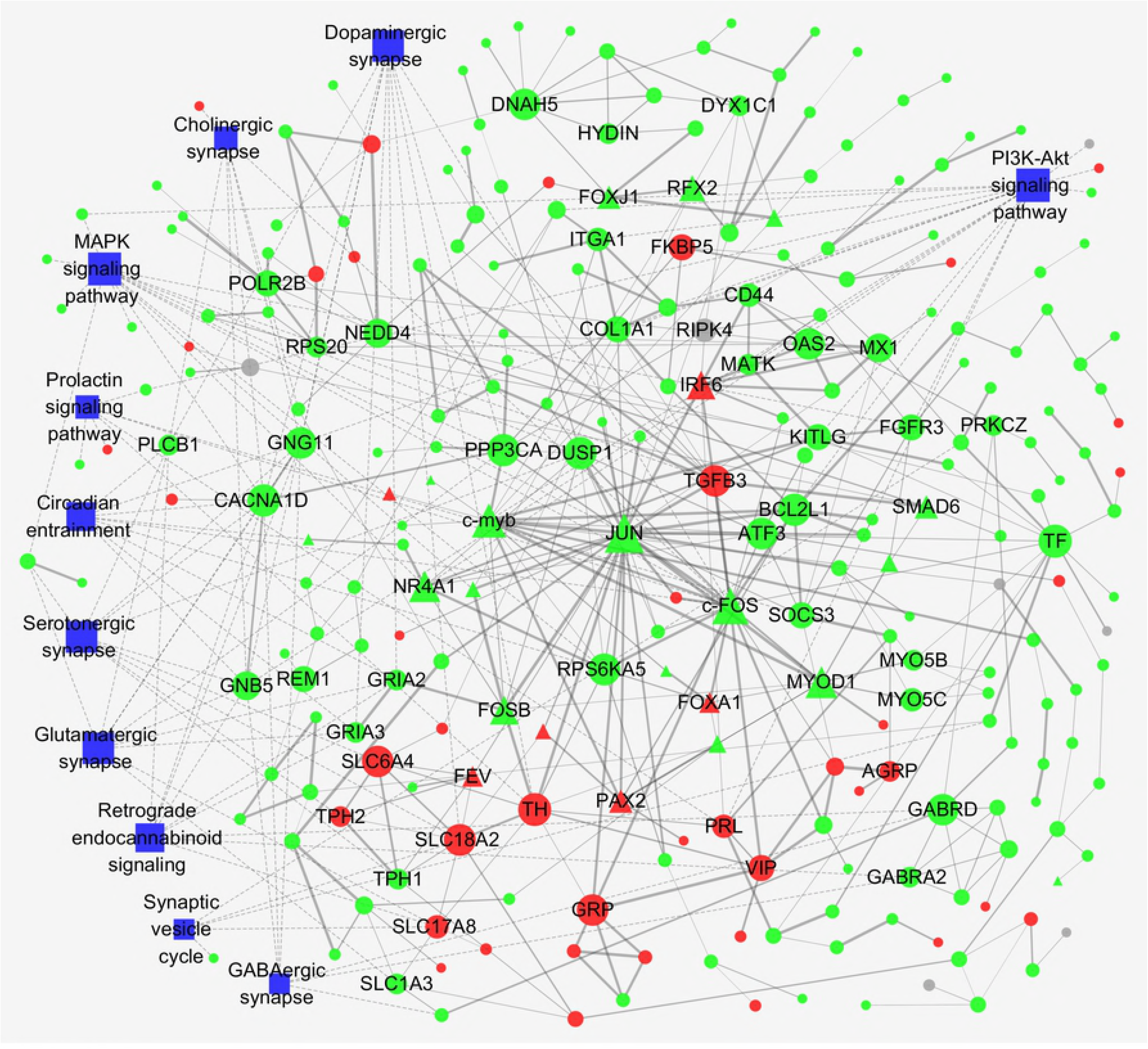
**Protein-protein interaction (PPI) networks of DEGs**. Circle nodes for genes/proteins, pentagon for transcription factors, rectangle nodes for KEGG pathway or biological process. Nodes with a degree of connectivity greater than 5 were labeled and all nodes in S1 Fig were labeled. According to trend analysis, genes/proteins were colored in red (Cluster 1, representing up-regulation over time), green (Cluster 1, representing down-regulation over time) and gray (Cluster 3, gene expression changed in BP stage only). Interaction were show as solid lines between genes/proteins, edges of KEGG pathway/biological process in dashed lines.

## Discussion

Reproduction is a complex and precisely regulated physiological process that requires the coordination of hypothalamus-pituitary-gonadal (HPG) axis for successful procreation. In this study, a new insight into Muscovy duck (*Cairina moschata*) hypothalamus transcriptome profiles in three reproductive Stages of egg-laying anaphase, broody prophase and broody metaphase were provided through deep sequencing. We selected the experimental individuals for transcriptome study according to behavioral observation and anatomical feature of the ovary.

One possibility is that oxidative stress might play a role in mediating reproductive costs during incubation[25,26]. We performed studies of GO enrichment of DEGs in pairwise comparisons of LA-vs-BP, LA-vs-BM and BP-vs-BM. The results showed that biological process of ‘response to oxygen levels’ and ‘response to oxygen-containing compound’ were both involved in the top 20 significantly enriched biological processes in three libraries (Fig 2III, S7 Table). A set of genes involved in the biological process, including *SOD3, NOS2, TH, TPH2* and *SESN1*, which response to oxidative stress or participate in the oxidation process. It suggested a possible role of oxidative stress might invoke reproductive costs that potentially change genes expression. Our study provided stronger evidence for the hypothesis that broodiness might cause oxidative stress in hypothalamus.

The control of avian reproduction involves the interaction of external stimuli with neuroendocrine[27]. It has proved the hypothalamic dopamine (DA), serotonin (5-HT) and Vasoactive intestinal peptide (VIP) up-regulated in the avian brooding period[28,29]. Results of KEGG analysis were consisted with previous study that many DEGs significantly enriched in Dopaminergic synapse and Serotonergic synapse pathway (Fig 2IV), and VIP gene up-regulated in brooding stage. In Dopaminergic synapse pathway, TH gene was up-regulated which encoded tyrosine hydroxylase enzyme. The product of the enzymatic reaction, L-DOPA, is a precursor to dopamine, has been shown to be the rate limiting step in the production of dopamine. In Serotonergic synapse pathway, *TPH2* gene was up-regulated. The translation protein of *TPH2*, Tryptophan hydroxylase, is an enzyme involved in the synthesis of the neurotransmitter serotonin. *SLC18A2* up-regulated in broody stage, this gene encodes a vesicular monoamine transporters protein (VMAT2), which transport several types of monoamine neurotransmitters, including dopamine and serotonin[30]. We found the synthesis and transport of DA and 5-HT are both elevated in the process from egg-laying to brooding.

In this study, glutamatergic synapse and GABAergic synapse pathway were significantly changed in the transition of reproductive cycle. The genes of glutamate receptor (*GRIA2* and *GRIA3*) and GABA receptor GABA_A_ (*GABRD, GABRA2*) were down-regulated in hypothalamus of broody duck. To date, studies showed that GABA, GLU and their receptors play special roles in mating and maternal behavior of rats [31,32]. Besides, it was reported that neurotransmitter of GABA and NPY stimulates food intake in chicks by interaction with the GABAergic system via GABA_A_ receptors[33,34]. The reduction of GABA_A_ may be critical reason for duck bad appetite during brooding. Our results first indicated glutamate and GABA signaling pathway also play critical roles in poultry reproductive behavior of broodiness.

Further analysis of genes expression trend and protein-protein interaction (PPI) networks we focused on 6 extra hub DEGs, including *GRP* (Gastrin-releasing peptide)*, FKBP5* (immunophilin)*, TTR* (transthyretin), DIO2 (type 2 deiodinase) and transcription factors of c-Jun and c-Fos. To date, it has been reported that GRP plays significant roles in many physiological processes, including food intake, circadian rhythms, male sexual behavior, and fear memory consolidation, through the specific GPCR, GRPR-mediated mechanisms[35]. As previously noted, the glucocorticoid receptor (GR) is a central receptor and a transcription factor in regulating the HPA axis. In fact, HPA axis dysregulation and GR resistance can be governed by changes in FK506-binding protein 5 (FKBP5) function[36]， and broody duck have increased expression levels of FKBP5 compared with egg-laying individuals in our study. Recent molecular analyses have revealed that thyroid hormone in the hypothalamus plays a key role in signaling day-length changes to the brain and thus triggers seasonal breeding in both birds and mammals[37]. Hypothalamic expression of deiodinase enzymes that catabolize thyroid hormone (T4) into the receptor-active triiodothyronine (T3) or T4-binding proteins (transthyretin) are strikingly regulated by changes in photoperiod and function as master regulators of seasonal reproduction in birds [38–40]. The genes encoding type 2 deiodinase (*DIO2*) and transthyretin (*TTR*) prominently down-regulated in the duck hypothalamus during broody stage, providing a rhythmic gating mechanism for thyroid hormone signaling. Transcription factors of c-FOS and c-Jun are components of activator protein-1 (AP-1) and play crucial roles in the regulation of diverse extracellular stimuli, which include neurotransmitters, pro-inflammatory cytokines, oxidative and other forms of cellular stress[41,42].

In conclusion, our results suggest that modified gene expression in the hypothalamus may account for the transition between the laying and brooding phases. We identified differential expression genes among three reproductive stages involved in oxidative stress. At the transcriptional level, characteristic changes included changes of neurotransmitters and related receptors during broodiness. The interactions of glucocorticoid and thyroid hormone signaling with regulator gene function as critical regulators of timing Muscovy duck reproduction. This study therefore advances our understanding of the biological processes regulated by reproductive cycle in hypothalamus.

## Acknowledgments

Special thanks to Anqing Yongqiang Agricultural Science and Technology Stock Co., Ltd. for the raising of Muscovy duck and collecting of phenotypic data due to which I am able to study the reproductive behavior.

## Supporting information

S1 Fig. Protein-protein interaction (PPI) networks of DEGs which all nodes labeled.

S1 Table. Mapped statistical results of 12 libraries

S2 Table. The differently expressed genes detected between LA and BP.

S3 Table. The differently expressed genes detected between LA and BM.

S4 Table. The differently expressed genes detected between BP and BM.

S5 Table. Differently expressed transcription factors.

S6 Table. Primer sequences used for quantitative PCR validation.

S7 Table. Lists of GO enrichment analysis of specifically expressed genes.

S8 Table. Lists of KEGG pathway analysis of specifically expressed genes.

